# pycoMeth: A toolbox for differential methylation testing from Nanopore methylation calls

**DOI:** 10.1101/2022.02.16.480699

**Authors:** Rene Snajder, Adrien Leger, Oliver Stegle, Marc Jan Bonder

## Abstract

Advances in base and methylation calling of Oxford Nanopore Technologies (ONT) sequencing data have opened up the possibility for joint profiling of genomic and epigenetic variation on the same long reads. Existing data storage and analysis frameworks that were developed for CpG-methylation arrays or short-read bisulfite sequencing data have severe shortcomings for handling of ONT data, failing to fully exploit methylation profiles obtained from long read technologies. To address these issues, we present **pycoMeth**, a toolbox to store, manage and analyse DNA methylation data obtained from long-read ONT sequencing data. Our toolbox centers around a new storage format called **MetH5**, which allows simultaneously for efficient storage of and rapid data access for read-level and reference-anchored methylation call data. Building on this storage format, we propose efficient algorithms for the segmentation and differential methylation testing of methylation calls from ONT data. Our methods draw from read-group and read-level information, as well as methylation call uncertainties, and allow for *de novo* discovery of methylation patterns and differentially methylated regions in a haplotyped multi-sample setting. We show that **MetH5** is more efficient than existing solutions for storing ONT methylation calls, and carry out benchmarking for segmentation and differential methylation analysis, demonstrating increased performance and sensitivity of pycoMeth compared to existing solutions.

## Background

High-throughput technologies for profiling of DNA base modifications have become established tools to study epigenetic regulation. In mammalian cells, the predominant and most studied type of base modification is the methylation of cytosine in the 5’CpG3’ context (often abbreviated 5mC or simply CpG-methylation). Genomic regions enriched with this CpG motif (often referred to as CpG-islands, CGI) are found to be less tightly associated with nucleosomes, hence more accessible to DNA-binding proteins such as transcription factors [1]. Methylation of CpG in regulatory regions can affect gene expression in a variety of ways, including via direct interference with transcription factor binding or via recruitment of binding proteins attracted to methylated CpGs [1, 2]. Other, arguably less well studied, types of DNA base modifications include the methylation of adenine in 5’GATC3’ (6mA) context or any of the oxidative derivatives of 5mC (5hmC, 5fC, and 5caC) [3].

There exists a growing repertoire of high-throughput assays for the profiling of CpG-methylation states. Aside from targeted approaches like bead array based methods, the most commonly used techniques are based on short read sequencing and offer single-base resolution and genome-wide coverage [4]. These include whole genome bisulfite sequencing (WGBS), and more recently an enzymatic methylation sequencing protocol, promising lower DNA degradation and more balanced base representation [5]. Both of these methods have been applied in bulk and single cell setting.

Our increased understanding on genetic variation has created demand for longer reads, and has therefore given rise to long-read sequencing technologies such as those developed by Pacific Biosciences (PacBio) as well as Oxford Nanopore Technologies (ONT), which can directly sequence native DNA molecules. Sequencing long single molecules can aide in problems that are difficult to resolve with short-read sequencing, such as the reliable detection of structural variations [6], phasing of variants into paternal and maternal haplotype [7], as well as the assembly of an entire human genome including low-complexity regions [8]. Additionally, ONT sequencing datasets can be (re)processed to obtain measurements of base modification states [9]. This allows for a host of applications through profiling the epigenome in a haplotype-resolved, whole-genome, single-molecule setting [10], including in regions which are poorly studied and annotated.

These new opportunities come with new challenges that need to be addressed in downstream analysis software. In the initial step, the modification state of a base needs to be determined. For this, a number of methylation callers have already been published, including Nanopolish [10], DeepSignal [11] and ONT’s own Megalodon [12]. These methods have been compared and benchmarked elsewhere [13]. The first pivotal challenge which arises from this new data type, and one which we address, is the need to efficiently store and retrieve base-level information with read association. We focus specifically on methylation calls from Nanopolish, a Bayesian methylation prediction tool emitting log-likelihood ratios (LLRs) of methylation [10]. The large number of methylation calls (up to 850 million CpG-methylation calls in a 30x coverage sequencing experiment of the human genome [14]) need to be stored and made accessible in an efficient manner. HTSLib [15] recently implemented two new tags (MM and ML) to store base modification probabilities together with the read alignments in the SAM format, storing methylation calls efficiently but at the cost of access speed (**Figure 2C**). Here, we propose **MetH5**, an open standard storage format optimized for rapid random access, scalability, and parallel computing, storing individual methylation calls at read and base level.

Furthermore, standard downstream analysis tasks such as discovery of differential methylation require software tools supporting methylation data from ONT sequencing. While existing methods designed for methylation frequency analysis measured using bisulfite sequencing can be appropriated, this approach ignores read information and requires binarizing methylation probabilities to discrete methylation calls. Downstream analysis software which takes full advantage of the probabilistic nature of ONT methylation calls as well as the molecule-level information, such as haplotype assignment, is currently lacking. To close this gap, we provide **pycoMeth**, a software suite for *de novo* methylation segmentation and differential methylation testing. To shed dependence on reliable genome annotation and to identify methylation patterns *de novo*, pycoMeth implements a Bayesian changepoint detection framework as a methylome segmentation tool. This method takes into account read-level information, as well as methylation call uncertainties, and produces a consensus segmentation over multiple read-groups (e.g., multiple samples, haplotypes, or sample/haplotype combinations). Furthermore, pycoMeth’s differential methylation testing module offers a variety of testing options for the discovery of differentially methylated regions (DMRs) in two or more samples, as well as a reporting function generating HTML reports for discovered DMRs.

Together, in this work, we present a software toolbox that represents an accessible, integrated solution, addressing the unique challenges of methylation analysis from ONT sequencing, encompassing efficient storage, segmentation, and differential methylation testing (**Figure 1**). To showcase our software, we also perform an extensive benchmark on simulated and real data, demonstrating the efficiency, flexibility, and versatility of **pycoMeth** and the **MetH5** format compared to existing tools for ONT and bisulfite sequencing.

**Figure 1:**
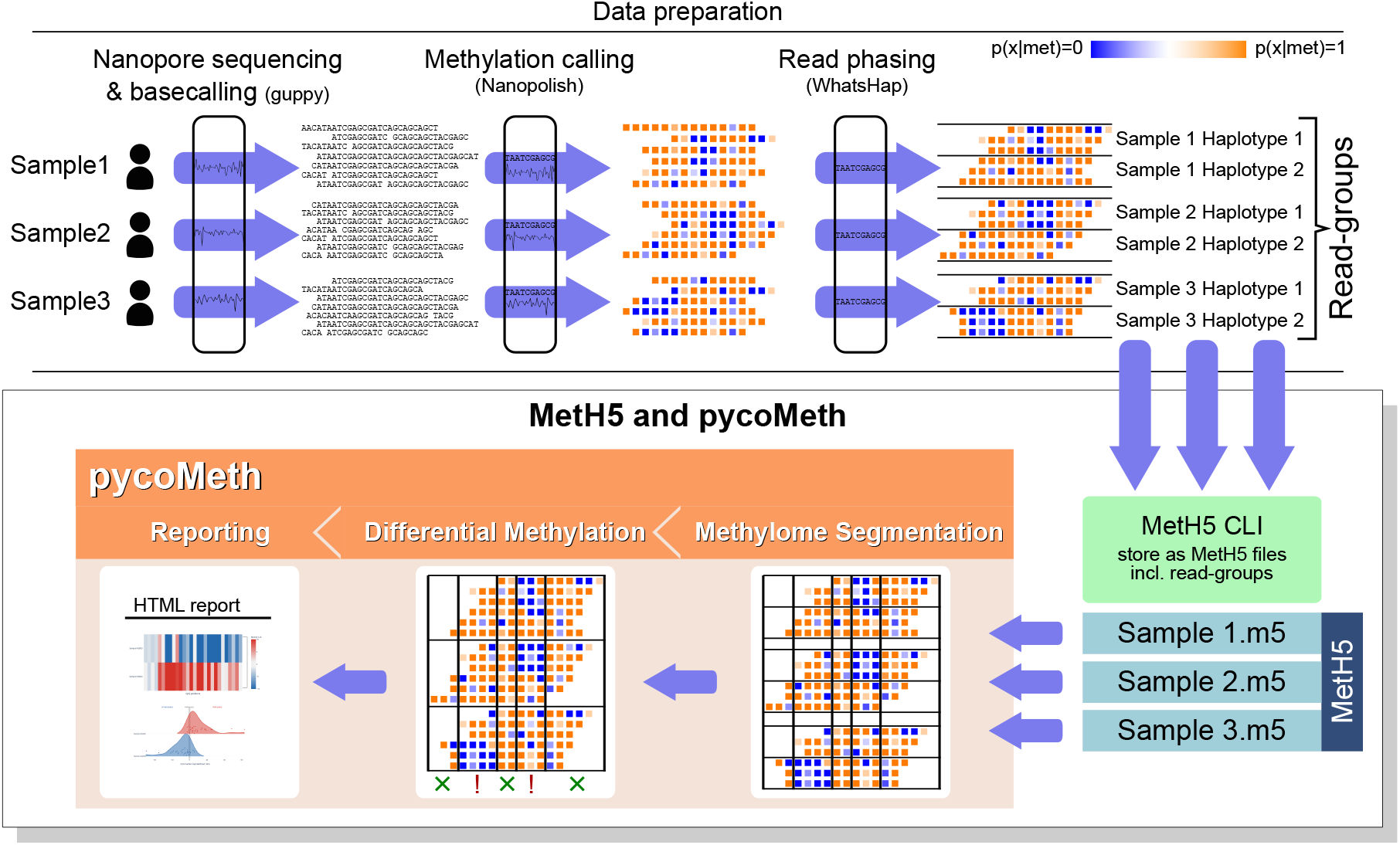
Overview of differential methylation analysis on ONT data using MetH5 and pycoMeth. Here, three biological samples are ONT-sequenced, base- and methylation-called, and haplotyped. The methylation caller output is stored in MetH5 format. Sample name and haplotype assignment for each read are stored as read groups in the MetH5 file (**Figure 2**). PycoMeth methylome segmentation is performed, producing a consensus segmentation over all read-groups. Each segment is then tested for differential methylation or allele-specific methylation (in this example between-sample differences, ignoring haplotypes). Finally, pycoMeth generates a summary HTML reports as well as detailed reports for all DMRs (**Figure 3**).

## Results

### MetH5 - an efficient read-level base modification container

Data storage and retrieval in pycoMeth is handled by MetH5 - a file format specifically designed for base methylation call storage from long read data with uncertainties (such as ONT). MetH5 enables rapid random access and is optimized for parallel computing, while retaining access to all long-read specific information such as methylation call uncertainties and molecule-level information. The MetH5 format (**Figure 2A**) is built on the Hierarchical Data Format (HDF) version 5[16], and we consider the following guiding principles for its design. *Read-level storage*: All base modification calls are stored together with the read they originated from, in order to allow read-level and read-group-level analyses. *Base-level uncertainty estimates*: MetH5 is not limited to binary calls, but can natively store the confidence values output by the base modification caller. *Rapid random access*: Base modification calls are stored in order of their genomic coordinate and indexed such that they can be retrieved with minimum disk I/O (**Figure 2B-C**). *Parallel processing*: Chunked storage and accessor methods facilitate even load distribution when used in parallel systems (**Supplementary Figure 1**). *Efficient storage*: Using efficient data types, data compression, and avoiding data duplication (such as read names or chromosome names). *Flexible annotations*: Reads can be annotated with arbitrary read-group qualifiers (e.g. sample, haplotype group, haplotype id). We evaluate the runtime performance and storage efficiency of the MetH5 format in **Figure 2C**.

**Figure 2:**
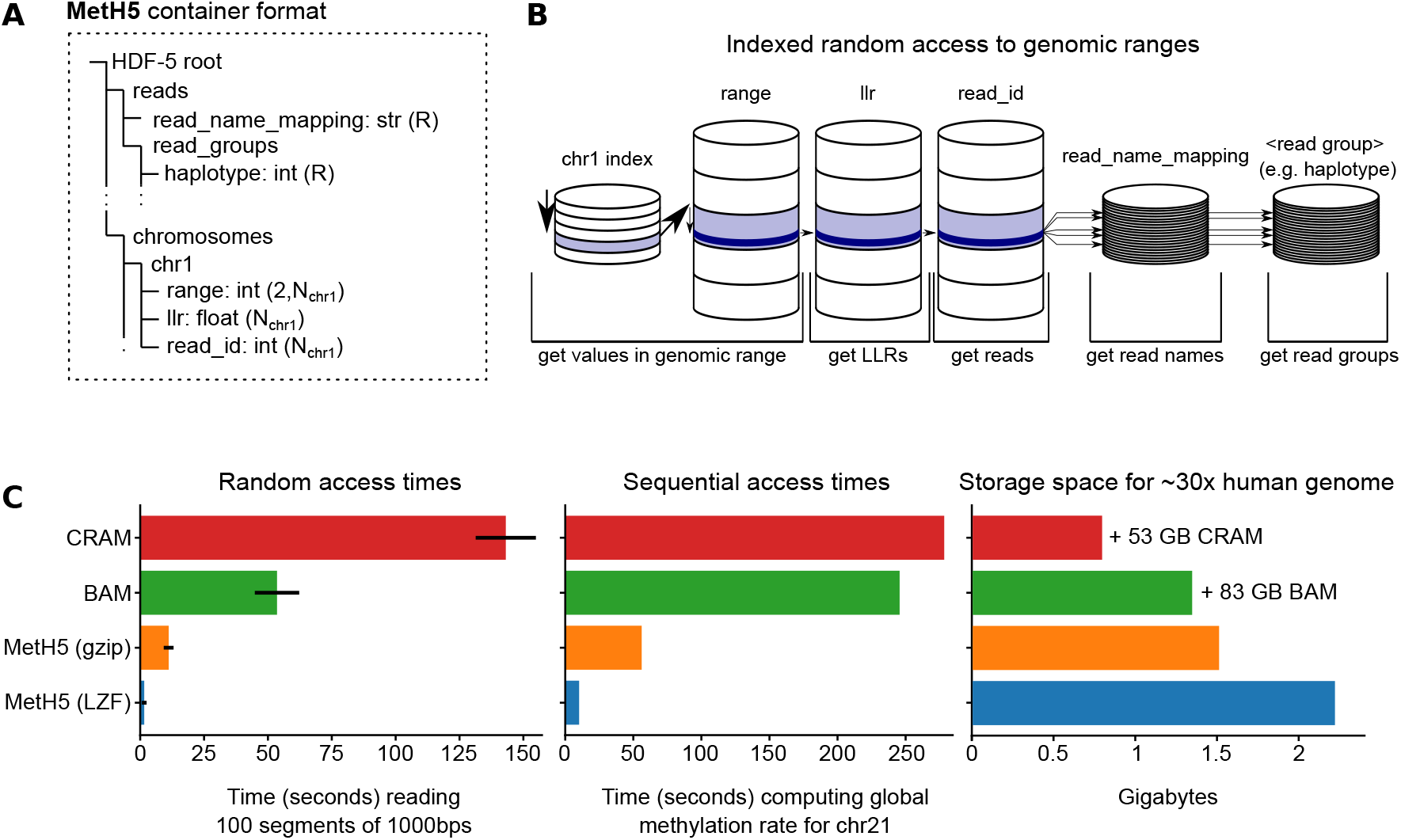
The MetH5 file format. **A)** Structure of the HDF5 container including dataset types and shapes. *N*_*x*_ refers to the number of methylation calls per chromosome *x. R* refers to the total number of reads in the entire container. Methylation calls are stored together with their genomic coordinate on the chromosome (range), the log-likelihood ratio (LLR) of methylation, and a numeric read ID (unique to this container). Read names are optionally stored, mapping each of the MetH5 numeric read IDs to the original read name. An arbitrary number of read groupings can be stored, assigning each read to exactly one read group per grouping. **B)** Schematic representation of random access in the MetH5 format. An index per chromosome allows direct access to the required chunk. The range dataset can then be searched for the start and end index. Once these indices have been acquired, LLRs and read IDs can be read directly and optionally. If globally unique read names are required, they can be looked up directly using the read ID, and the same holds for read groups such as haplotype assignments. **C)** Performance comparison between MetH5 and BAM/CRAM format with MM tag (**Methods**). In the file size comparison, bars represent only the extra space occupied by MM and ML tags, and native BAM size is annotated next to the bar.

MetH5 comes with a python API to abstract the architecture and provide developers with a coherent software interface. In addition to the python API, the meth5 package also comes with a command line user interface (CLI), which allows the creation, merging, and annotation of MetH5 files. The software also supports the extraction of key data quality statistics, such as base modification rates and coverage, for visualization in external software such as the integrative genomics viewer (IGV) [17, 18].

### pycoMeth Meth_Seg - Bayesian Methylome Segmentation for Haplotype-Aware Multi-Sample Changepoint Discovery

The ability to measure methylation on haplotyped long reads offers a unique opportunity for discovery of methylation patterns in a *de novo* fashion, independent of pre-made functional annotations or CGIs. Utilizing the efficient access to read-level methylation information offered by the MetH5 format, we implement pycoMeth Meth_Seg, a Bayesian changepoint detection algorithm (**Figure 3A**) for multi-read-group segmentation of methylation profiles, designed for the *de novo* discovery of methylation patterns from multiple (haplotyped) ONT sequenced samples. In contrast to previous segmentation methods, which either segment a single methylation profile [19], or derive a segmentation from differential methylation between two samples [20], pycoMeth Meth_Seg takes into account an arbitrary number of read groups (e.g., biological samples, haplotypes, or individual molecules/reads) to detect a dynamic set of methylation patterns from which it then derives a single consensus segmentation. To do so, the read-group annotation stored in the MetH5 container can be used to inform pycoMeth Meth_Seg about categorical methylation confounders such as biological sample or haplotype, which are then considered equally in the segmentation, allowing for for haplotype aware multi-sample methylome segmentation.

**Figure 3:**
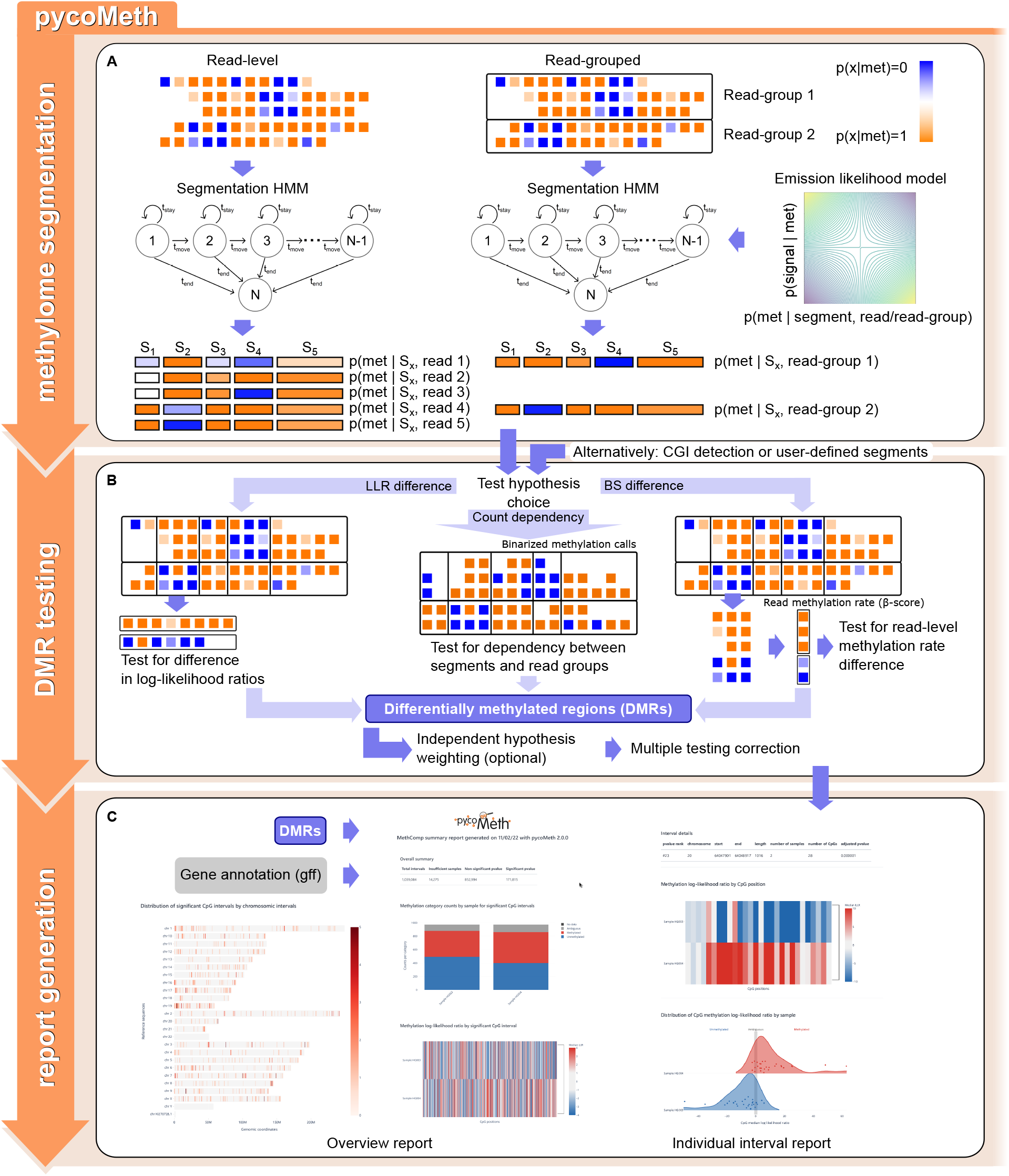
Example pycoMeth workflow for differential methylation analysis. **A)** Methylome segmentation using a Bayesian changepoint detection model. Segmentation can be computed on a read-group (e.g., haplotype) level. Emission likelihood in the HMM models methylation call uncertainties as well as an optional methylation rate prior. **B)** Differential methylation testing allows for a number of test choices. The LLR difference hypothesis compares methylation call LLRs within a segment between two samples directly. Selecting the count dependency hypothesis or the *β*-score difference hypothesis (default) both result in binarization of methylation calls based on a defined LLR threshold. The count dependency hypothesis leads to a test on contingency tables, testing dependency between methylation count and read group, whereas the *β*-score difference hypothesis results in a test comparing, for each segment, the read methylation rates between read groups. Regardless of test hypothesis, p-values are then subjected to multiple testing correction. **C)** The reporting module generates an overview HTML report, as well as individual interval reports.

The pycoMeth Meth_Seg model directly models methylation calls as uncertain observations, based on the LLRs stored in the MetH5 container. Methylation rates per segment per read group are then estimated as the parameters of the model, while simultaneously the segmentation is optimized to maximize the likelihood of observations (**Methods**). Default hyperparameters are set to maximize sensitivity such that typically an oversegmentation is achieved. This results in a segmentation that is unbiased towards between-sample differential methylation, as it is designed to compute a segmentation even in the absence of DMRs.

Segmentation can be performed either via a python API, or using a CLI which takes one or more MetH5 files as the input. PycoMeth can take advantage of MetH5’s chunked data storage, allowing chunked operations in order to allow efficient load distribution on parallel systems (**Supplementary Figure 1**).

### pycoMeth Meth_Comp - Versatile Differential Methylation Testing Suite

Once the methylome has been segmented, pycoMeth can be used to test segments for differential methylation between two or more samples, implemented in pycoMeth’s Meth_Comp subcommand. Methylation calls can either be provided as one MetH5 file per sample, or a single MetH5 file containing read-group annotations, such as when testing for allele specific methylation (ASM) within a single sample using read groups to tag haplotypes. PycoMeth Meth_Comp accepts any region information, provided in bed-format, for differential methylation testing (**Figure 3B**), and also implements a heuristic CGI detector (pycoMeth CGI_Finder) for the study of CGI methylation.

A variety of differential methylation test options are provided by pycoMeth Meth_Comp. The user can define a test hypothesis, and pycoMeth chooses a test depending on the number of samples, the test hypothesis, and other parameters (**Methods**). For multiple testing correction, pycoMeth implements a number of options for p-value adjustment, including an optional independent hypothesis weighting (IHW) scheme [21] that uses standard deviation of methylation rates as a weight in order to draw more power from a large number of tests. If more than two samples are provided, the matching 2-sample test will be performed as a *post-hoc* test in a one-vs-all setting for each interval where the null hypothesis had been rejected in the n-sample test. DMRs are reported as a tab-separated file. The optional Comp_Report subcommand provides additional functionality to generate easily accessible HTML reports, including visualizations in the context of user provided gene annotations (**Figure 3C**).

### Benchmarking pycoMeth

In order to evaluate performance of the different tools in the pycoMeth suite, we applied pycoMeth to two datasets. The first is a simulated Nanopolish dataset of methylation calls on chromosome 1(hg19) on two samples with differential methylation between them. We started by simulating methylation patterns organized in segments with either high, low, or intermittent methylation rate, with about 8% of the segments differentially methylated between the two samples with a variety of effect sizes (from 0.15 to 0.6 difference). This information was used to generated a low-coverage (15x) and a high-coverage (30x) dataset by simulating reads and LLRs, approximating the uncertainty distribution of Nanopolish methylation calls (**Methods**). The second dataset is a real ONT sequencing dataset from a father-mother-son trio sequenced by the Genome in a Bottle (GIAB) consortium [22] (information on alignment and data processing can be found in the methods). Here as well, we investigated the effects of a high-coverage (30x per sample) and low-coverage (15x per sample) setting. This synergy between the simulated and the real datasets allows us to draw conclusions about the accuracy of our methods in the real data.

### MetH5 facilitates rapid random access to methylation calls

To show the benefits of the MetH5 container format, we compared our format with the current definition and implementation of modification scores in hts-specs (MM and ML tag, samtags repository version df69c35). Here, we focused only on the data from one of the GIAB samples (HG003, Methods), either using BAM files combined with the MM tag or stored using the MetH5 format (**Methods**). Methylation calls were stored in MetH5 as well as BAM and CRAM format. Assessing the advantages of the MetH5 file format, **Figure 2D** compares random access and sequential access times of methylation scores stored in MetH5 versus CRAM and compressed BAM format, as well as storage space required. While BAM/CRAM files make more efficient use of storage, between 165 to 1,400MB (11 to 64% of that used by MetH5) depending on compression, they are significantly more expensive to read (about 5 to 90 times slower depending on compression).

### pycoMeth changepoint detection on simulated data outperforms existing tools

Next, we assessed the performance of the segmentation. Due to the lack of tools designed for segmentation and differential methylation analysis in single-molecule sequencing data, we instead compare our segmentation with two commonly used methods designed for segmentation of methylation rates from bisulfite sequencing data, methylKit [19] and MethCP [20]. MethCP performs segmentation of a differential methylation profile in a two-sample setting, which is why it is naturally biased towards DMRs. In order to also compare with an impartial segmentation, we use methylKit’s single-sample segmentation on the summed methylation profile of the two samples which are compared.

We created two segmentations using pycoMeth Meth_Seg, one with a maximum of 16 segments per 300 CpG sites and one with the same maximum per 600 CpG sites (which we refer to as “pycoMeth coarse”). Additionally, we evaluated the performance of pycoMeth Meth_Comp when matching the settings to those used by methylKit and MethCP, in order to investigate the impact of the segmentation and DMR testing methods independently.

To evaluate the segmentation quality, we counted the number of ground-truth segments whose changepoints (segment boundaries) are represented in the segmentation in the high-coverage example (**Figure 4A**) and the low-coverage example (**Supplementary Figure 5A**). As an accuracy threshold, we define that a changepoint has been correctly identified if the nearest predicted changepoint is no further away than 5% of the containing segment’s width. The pycoMeth segmentation identified 72.2% of all DMR segments and 72.9% of non-DMR segments (full match). For 24.1% of DMR segments and 23.5% of non-DMR segments, one side of the segment (start or endpoint) could be accurately detected (partial match). Only 3.6% of DMR segments and 3.6% of non-DMR segments could not be accurately detected on either side. However, the segmentation also predicted 33,100 additional changepoints which do not correspond to ground-truth segments (+485% oversegmentation). Reducing the segmentation granularity (pycoMeth coarse) reduced the oversegmentation to 18,276 (+268%) additional changepoints while reducing detection power by about 4%. The methylKit segmentation, which resulted in the least number of segments (only 35% oversegmentation), still captured a fair number of DMR (33.5% found, 32.2$ partially found) and non-DMR (50.7% found, 30.4% partially found) segments, whereas MethCP showed good performance on DMR segments (63.4% found, 22.4% partially found), but failed to capture non-DMR segment boundaries (15.4% found, 28.7% partially found) despite producing a large number of segments (667% oversegmentation). Since detection power naturally also increases with segmentation granularity, we performed a randomization test in which predicted segments are shuffled and compared to the original segmentations, confirming that all segmentation methods perform vastly better than random 3.

**Figure 4:**
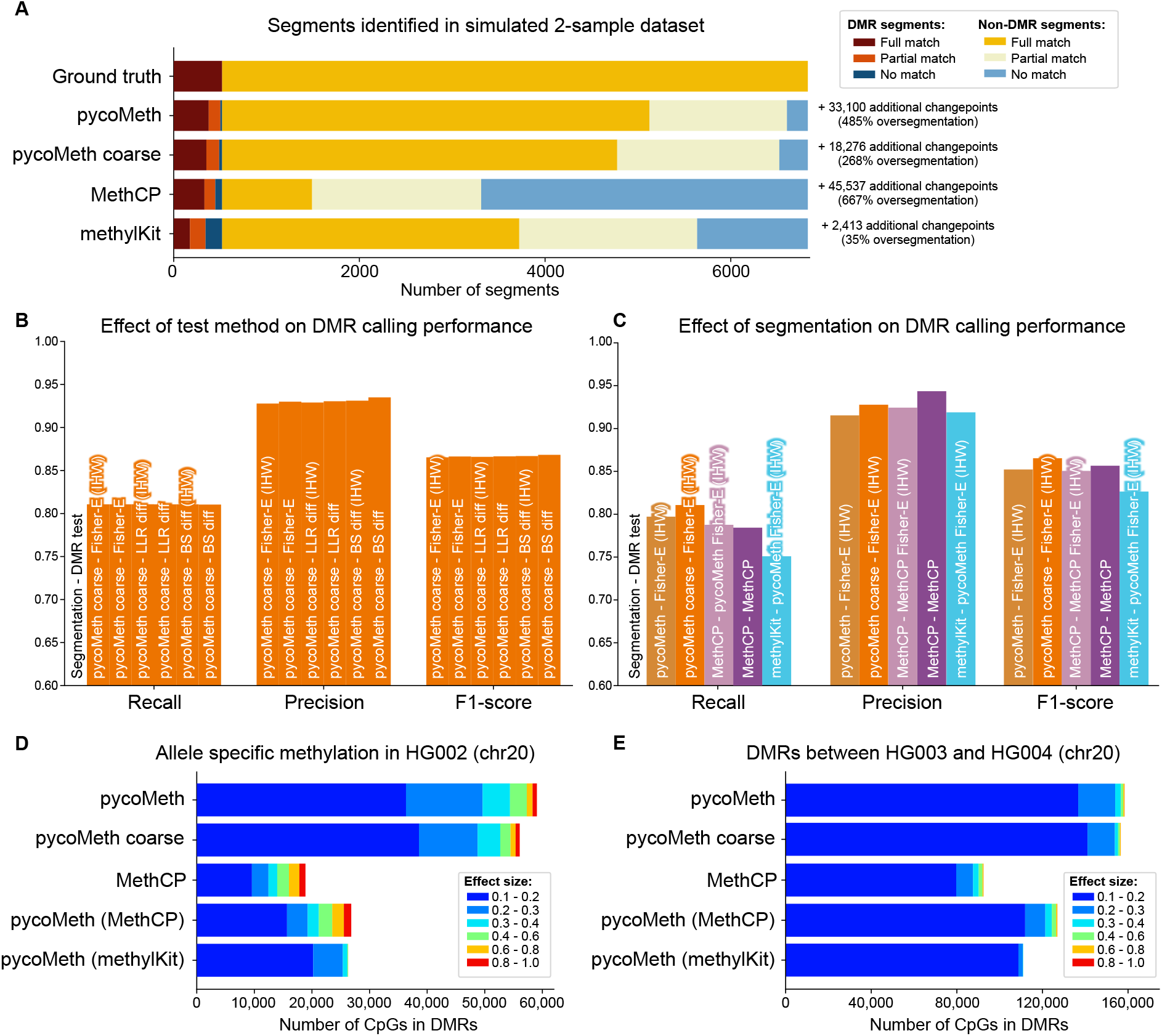
Benchmark on simulated data and GIAB dataset. **A)** Number of segments identified as full match (both the true start and end changepoint have been accurately identified), partial match (one of either the true start- or end-changepoint has been accurately identified), or no match (neither the true start nor end changepoint have been accurately identified) in the high-coverage simulated dataset. For the purpose of this graphic, a changepoint is considered accurately identified if a predicted segment breakpoint is no more than 5% of its containing segment’s length away from the true (ground truth) changepoint. Additional changepoints are the number of predicted changepoints not counting towards a partial or full match (oversegmentation). **B)** Precision, Recall, and F1-score computed on the simulated dataset for all test methods implemented in pycoMeth, tested on the pycoMeth coarse segmentation. **C)** Precision, Recall, and F1-score computed on the simulated dataset using the Fisher exact test, compared between different segmentation methods. **D)** Allele-specific methylation called on HG002. PycoMeth calls were performed using the Fisher exact test with IHW. Colors represent the effect size of the containing segment (absolute differential methylation rate). **E)** DMRs identified between HG003 and HG004.

After segment identification, we performed DMR testing using pycoMeth Meth_Comp on all four segmentations. First, we assessed the impact of different statistical tests on DMR identification in the high-coverage example, comparing the options implemented in pycoMeth with MethCP **Figure 4B**. We observe that in this setting, the test method used in pycoMeth Meth_Comp does not strongly impact the outcome, with the improved precision of the most conservative test (*β*-score difference) leading to the best overall performance (F1 score 0.868, used to represent the compromise between precision and recall) compared to the LLR difference test and Fisher exact test (both with an F1 score of 0.866). Next we tested the impact of the segmentation on DMR identification, **Figure 4C**, fixing the test to Fisher exact (count dependency hypothesis) - as this is the same test used in MethCP - with IHW. We find that both pycoMeth options have higher recall than the other tools. While MethCP shows best precision, pycoMeth coarse has the highest F1-score. In summary, of all configurations, pycoMeth coarse with *β*-score difference test yielded the best performance in terms of F1-score and has been set as the default parameters when using pycoMeth.

In the low-coverage (15x) example with the same test settings (**Supplementary Figure 5B-C**) we found that reduced coverage led to a slight reduction in performance in all methods. Most affected was methylKit, with an F1 score reduction of 0.036, compared to MethCP (0.016 reduction) and pycoMeth coarse segmentation (0.013 reduction). Next, we investigated the agreement of changepoint predictions across low- and high-coverage examples. The Jaccard index of changepoints found is 0.83 in the pycoMeth segmentation, 0.82 in pycoMeth coarse, 0.85 in MethCP and 0.76 in methylKit, indicating good stability in all methods.

### pycoMeth shows high power at detecting low-effect DMRs in real ONT data

Besides simulations, we also assessed pycoMeth Meth_Seg performance in a real-world setting, by generating a methylome segmentation on the GIAB data, and test both for DMRs between HG003 and HG004 (parental samples), as well as trying to identify ASM in HG002 (son) **Figure 4D-E**. For the between-sample DMR test, coverage was 30x per sample, and in the ASM test 15x coverage per haplotype, analogous to the high- and the low-coverage simulation examples, respectively. We find that pycoMeth identifies a much larger number of CpGs in DMRs, particularly in DMRs with low effect sizes. Examining chromosome 20 as a benchmark, pycoMeth coarse with the Fisher exact test and IHW identifies 77,054 DMR CpGs in the ASM scenario, where 66,307 (86.1%) were of low effect size (0.1 to 0.3) and 10,747 (13.9%) of high effect size (> 0.3). Using the same settings, in the parent comparison, pycoMeth identified 156,584 CpGs in DMRs, of which 153,839 (98.2%) were of low effect size and 2,746 (1.8%) of high effect size. Comparable figures for MethCP show lower numbers overall, but with more CpGs from high-effect-size DMRs: in ASM, 42,936 CpGs total with 29,450 (68.6%) of low effect size and 13,486 (31.4%) of high effect-size, and in the parent comparison, 92,435 total with 87,586 (94.8%) of low effect size and 4849 (5.2%) of high effect-size. Using pycoMeth’s Fisher-exact IHW test on the methylKit segmentation shows lower counts overall, with very few CpGs in high-effect-size regions: in ASM, 33,826 CpGs total with 31,591 (93.4%) of low effect size and 2,235 (6.6%) of high effect size, and in the parent comparison, 110,975 total with 110,853 (99.9%) of low effect size and 112 (0.1%) of high effect size.

We also find that the segmentation has a great impact on which segments are found, implying that the methods are complementary. Looking again at chromosome 20 and comparing with the tests on simulated data, only 14.8% of CpGs in DMRs found in the parent comparison are found by all segmentations, whereas in the simulated case, the agreement was 52.6%. Even between pycoMeth and pycoMeth coarse, there was only a 52.7% agreement. Comparing different tests on the same pycoMeth segmentation, 79.5% of sites are called by all tests, with the more conservative *β*-score difference method standing out, not calling 12.2% of sites which are called by the other two methods. On the simulated data, consistency is better, with 52.6% agreement across tools and 96% agreement across pycoMeth tests (**Supplementary Figure 6**).

Finally, seeing how pycoMeth has high power in low-effect-size settings, we evaluated overcalling of low-effect-size DMRs in both simulated and real data. On GIAB data, a randomization test was performed, in which LLRs in the MetH5 containers of HG003 and HG004 were randomized within each chromosome, such that methylation becomes fully independent of read, read group, and genomic location. A segmentation and DMR testing run between randomized HG003 and randomized HG004 was then performed to estimate a false discovery rate (FDR). We find this FDR is in the range of 0.07-2.2 for effect-size thresholds up to 0.1. For thresholds of 0.1 or higher, FDR ranges (mostly) from 0.04 to 0.23 for LLR-Diff and the Fisher exact test, and consistently stays below 0.05 for the BS-Diff test (**Supplementary Figure 7**). Furthermore, we find that the majority of false-positive segments are long (> 3,000 methylation calls per segment) low-effect-size DMRs which are effectively filtered out by the effect-size filter of 0.1. Computing precision and recall on simulated data at different effect-size thresholds also shows that segmentations with higher granularity suffer from reduced precision, with pycoMeth coarse segmentation resulting in the best precision and recall (**Supplementary Figure 4**). Taking everything together, pycoMeth coarse segmentation with the BS-Diff test hypothesis and IHW presents a good configuration for further studies of all ranges of methylation effects.

## Discussion

In this work, we presented pycoMeth and MetH5, a toolbox for the analysis of ONT-derived methylation data, encompassing storage, segmentation, and differential methylation analysis. We provided a benchmark on simulated and real sequencing data, each in a low- and high-coverage setting, comparing MetH5 and pycoMeth with existing solutions.

We observed that using MetH5 as a file storage showed a marked performance increase as compared to current BAM-based file formats for downstream methylation analyses, such as our segmentation. While we are aware of the advantages of having methylation calls stored in a well-supported file format, we believe MetH5 to be a complementary solution to methylation storage in BAM files, offering increased performance and allowing for easier sharing of methylation information due to being a compact specialized storage container, whereas BAM are large and monolithic, containing many different data modalities. On top of that, in our experiments, we currently observe compatibility issues between (mod)BAM files generated by Nanopolish and the modbampy and pysam libraries for downstream analysis, both with read-anchored as well as reference-anchored calls, but we expect that future improvements will solve this and potentially increase methylation accession speeds in BAM files. The use of the modification tags (MM & ML), is still new, and therefore support in downstream software is either not optimal or not existing, even though development is picking up speed rapidly. With MetH5, we provide a specialized file format allowing for extraction and storage of methylation data with the intent to simplify and speed up methylation-related analyses and methylation data sharing.

We have shown that the segmentation method used to determine segments for DMR testing has a great impact on the number of DMRs found in native data, particularly on DMRs with low effect sizes. Compared to existing DMR calling workflows, pycoMeth shows improved performance (F1 score) on simulated data, and identifies more DMR segments in real data. Furthermore, we show that the segmentation approach implemented in pycoMeth allows for the discovery of methylation changepoints, both in regions with and without DMRs, and demonstrates the potential for further study of *de novo* methylation pattern discovery from long-read sequencing in a multi-sample setting.

MetH5 and pycoMeth have been developed and tested with methylation calls from Nanopolish[9] in mind, as Nanopolish is easy to run on CPU-based hardware, has benefited from long-term maintenance and updates and has therefore become very robust and popular in the community. Furthermore, with Nanopolish being a Bayesian method, LLRs reported by it are well-suited for the uncertainty propagation method as implemented in pycoMeth Meth_Seg. However, newer base modification callers have shown higher methylation calling accuracy [13], and more recently, remora-based methylation calling has been integrated in the ONT basecallers guppy[23] and bonito[24], which store methylation calls as tags in BAM format. Future development of pycoMeth and the meth5 API will aim to support methylation calls in BAM format as the file standard and implementations stabilize, thus improving support for these methylation calling tools. Doing so will also enable us to investigate potential applications to methylation calls from PacBio sequencing, another avenue we intend to pursue in the future. Also, while mainly developed for the evaluation of CpG-methylation, all methods (aside from the CGI-finder) are also applicable to other types of epigenetic marks, such as adenine methylation, or cytosine methylation in GpC context.

## Conclusions

Here, we presented a toolkit and efficient file format for epigenetic analyses on ONT reads. Due to the novelty of single-molecule methylation calling, there are no gold standards yet for storage and analysis. With the MetH5 format, we attempt to provide an efficient method of storing reference-anchored methylation calls without compromising on read-level information or methylation call uncertainty information. The pycoMeth Bayesian segmentation method and differential methylation testing take advantage of read-level or read-group-level information, which tools designed for bisulfite sequencing typically do not consider. This new approach performs comparably to or better than previous tools in terms of segmentation accuracy and balance between DMR testing recall and precision (F1-score). Especially in a low-coverage setting and for the detection of methylation changes with low effect sizes, pycoMeth excels compared to the other tools tested.

## Materials and Methods

### GIAB Benchmark Data Preparation

GIAB raw fast5 files were downloaded from the Human Pangenome Project’s S3 bucket. In order to reach approximately 20x to 30x coverage, we use four flowcells from HG002 and three flowcells from HG003 and HG004, respectively (**Supplementary Table 1**). Phased SVs produced by [25] were downloaded from the NCBI ftp server (**Supplementary Table 1**). Reads have been re-basecalled using guppy version 5.0.11 with the high-accuracy model with modbases. Alignment to reference genome GRCh38 was performed using minimap2 [26] with the map-ont preset and otherwise default settings. Reads were haplotagged using whatshap[7] version 1.1. To produce Nanopolish[9] methylation calls in MetH5 format, we run Nanopolish call-methylation with Nanopolish version 0.13.3 and then use the python meth5 API to convert the Nanopolish output to the MetH5 format. In order to generate BAM files with MM tags, the “methylation_bam” branch (commit 9B01ad7) of Nanopolish has been used. BAM files were compressed to create CRAM files using `samtools view -C` and both BAM and CRAM were indexed using “samtools index”. Performance comparisons between MetH5 and BAM/CRAM files were performed using the meth5 version 0.8.0 and modbampy version 0.4.1 [27]

### Simulation of Methylation Profile and Nanopolish Methylation Calls

We use the tool OmicsSIMLA version 0.6 [28] with the parameter `--WGBS` to first generate a methylation profile based on human liver tissue for chromosome 1 (hg19) and simulate a control sample and a perturbed sample. We run OmicsSIMLA multiple times, with the `methy_theta` parameter (which indicates the effect size of DMRs) set to 0.15, 0.2, 0.25, 0.3, 0.35, 0.4, 0.45, 0.5, 0.55, and 0.6, each time with parameters `-p_diff_phase_meth 0.01 -p_diff_phase_unmeth 0.01 -p_diff_phase_fuzzy 0.01`. We combine the profile from these simulation runs, such that up to 10% of segments are DMRs. Segments which are highly methylated in control and up-methylated in the perturbed sample as well as segments which are unmethylated in control and de-methylated in the perturbed sample were treated these as non-DMR segments.

In order to then simulate Nanopolish methylation calls for these methylation profiles, we first estimate the distribution of read-lengths in a real ONT sequencing run as well as the distribution of LLRs in Nanopolish calls, based on the GIAB sample HG002. We model log10 read-lengths as a Gaussian mixture model with three kernels. Methylation call uncertainty encoded in LLRs from Nanopolish are modeled as a beta distribution.

Note that *p*(M|*X*), the probability of methylation given the observed raw signal *X* equals *σ*(LLR). We define certainty of a methylation call *c*(*LLR*) ∈ [0, 1] as *c*(*LLR*) = |*σ*(*LLR*) ∗ 2 − 1|, making it so that an LLR of 0 leads to *c* = 0, complete uncertainty of the methylation call, and *c*(*LLR*) = *c*(−*LLR*). We collect all certainties for chromosome 21 of HG002 and estimate their distribution as a beta distribution *B*(*α, β*). The resulting parameters of the distribution were *α* = 0.640308, *β* = 0.208756.

Methylation rate *µ* per CpG-site per sample were taken from the OmicsSIMLA simulations. We draw a random methylation status *x* ∼ B(*µ*) where *x* is either 0 or 1. Furthermore, we draw a methylation call certainty *c* ∼ *B*(*α, β*). The simulated LLR is then computed as LLR = *σ*^*−*1^(0.5 + (*x* − 0.5)*c*)

Nanopolish methylation calls were then simulated by randomly drawing read position and read length, followed by sampling methylation states for each CpG site covered by the read using the methylation rates simulated by OmicsSIMLA. Then the uncertainty distribution is used to sample an LLR for the methylation call. Both the high coverage and the low coverage dataset are drawn from the same methylation profile simulated by OmicsSIMLA, but reads were independently simulated. We simulate a total of 500,000 reads per sample for the high coverage dataset (corresponding to roughly 30X coverage) and 250,000 for the low coverage dataset (corresponding to roughly 15X coverage), and store the methylation calls in a MetH5 container using the meth5 python API.

### Implementation

All tools have been implemented in python and require python version 3.7. The MetH5 format implements an HDF version 5 [16] container which is accessed using the h5py[29] library. Other open source software libraries used in this work include NumPy [30], SciPy [31], pandas [32], pyfaidx [33], statsmodels [34], and plotly [35].

### Implementation of MetH5

The MetH5 format (**Figure 2A**), an HDF version 5 container, contains two top-level groups: chromosomes and reads. The chromosomes group contains one group named for each chromosome or contig, which in turn contain four datasets. The first three datasets, llr, read_id, and range, are all of length *n* and chunked using a chunk size defined upon container creation. They store the methylation call uncertainties (as a floating point number), locally unique read identifier (int) and genomic range (start and end integer coordinate on the chromosome, thus facilitating grouped methylation calls), respectively. Finally, upon indexing, another dataset chunk_ranges of dimension (*c*, 2), where 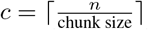 the number of chunks, is created. This dataset serves as an index for rapid random access of genomic coordinates. The second top-level group “reads” is entirely optional and stores read annotations. It contains the dataset read_name_mapping, a string dataset of shape *r*, where *r* is the total number of unique reads. This dataset stores read names and can be directly indexed using the local read identifiers stored in the read_id dataset. Additionally, the group “read_groups” can contain a variable number of datasets of shape *r* which can be used for read-group annotation, such as sample assignment or haplotype assignment. Users can define dataset compression options upon creation. As a default, lzf compression is enabled.

The meth5 python API and CLI implements creation of a meth5 file from Nanopolish result files. Random access to a genomic range is implemented by first identifying the required chunks inside a chromosome, using the chunk_ranges dataset, and then performing a binary search for the start and end index within the required chunk of the ranges dataset. These indexes can then be used to directly access the corresponding values in the read_id and llr datasets. Alternatively, chunk-based accessor functions allow direct access to the llr, read_id, and range datasets within a given chunk, and optionally also allow for inclusion of data from neighboring chunks whenever the methylation calls for a coordinate are split by the chunk boundary.

### Implementation of Bayesian Changepoint Detection

PycoMeth Meth_Seg implements a Bayesian changepoint detection algorithm modeled as an HMM, based on the segment based model defined in [36] and modified to account for variable number of segments by introducing a transition from any state to the end state, multiple read groups, and uncertainty propagation from methylation inference.

The number of states *S* is a hyperparameter and represents the maximum number of segments. Transition probabilities are defined as:

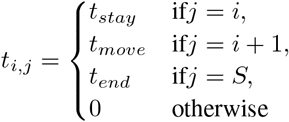

where *t*_*stay*_ = 0.1, *t*_*move*_ = 0.8, and *t*_*end*_ = 0.1 are priors controlling the granularity of the segmentation and their default parameters respectively.

Each segment is parameterized with *µ*_*s,g*_ ∈ [0, 1] the methylation rate of segment *s* in read-group *g*. If each read is represented as its own read-group, we refer to this as a read-level segmentation and *g* refers to the read. Emission likelihoods for each methylation call from segment *s* and read-group *g* given raw Nanopore signal *X* are derived as follows. Let *p*(*U* |*X*) and *p*(*M* |*X*) be the probability of a base being unmethylated or methylated respectively, given observed raw signal *X*.

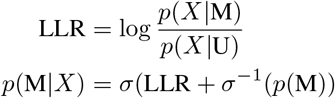

where *σ* refers to the sigmoid function 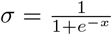 and *p*(M) = 1 − *p*(U) is the prior methylation probability.

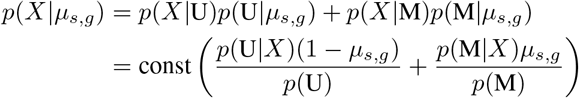

A segmentation is then computed using the Baum-Welch algorithm. Let *ψ* be the mapping between CpG sites and segments. In the expectation step we compute the posterior *p*(*ψ*(*i*) = *s*|*X, µ*) for all *i, s* using the forward-backward algorithm. In the maximization step, we then update the segment methylation rate parameter *µ* to the maximum likelihood estimator. Let *r*(*g*) be the set of all reads in read-group *g*.

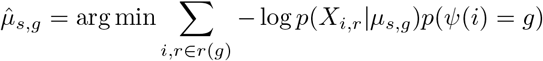

The expectation and maximization steps are repeated until all parameters *µ*_*s,g*_ have reached convergence with a tolerance of 1*e* − 4. Finally, a cleanup step is performed in which segments shorter than 5 CpG sites are merged with the next segment. To mitigate oversegmentation, neighboring segments whose methylation rate parameters differ less than 0.2 in all read-groups are then merged in a post-processing step. Since the memory requirement of the Baum-Welch algorithm scales *O*(*NSG*) where *N* is the number of CpG sites, *S* is the number of segments, and *G* is the number of read-groups, we perform the segmentation in a windowed fashion, on 300 or 600 CpG-sites per window by setting the corresponding parameter of pycoMeth Meth_Seg. This leads to artificial breakpoints between windows, causing some over-segmentation.

### Implementation of Differential Methylation Testing

DMR testing implements a number of different statistical tests depending on the test hypothesis, number of samples, and other parameters. Currently, three test hypotheses are implemented, named llr_diff, bs_diff, and count_dependency (**Supplementary Figure 2**). The test hypothesis llr_diff tests whether there is a significant difference in mean LLR between samples by computing a ranked test. This mode assumes all LLRs are independent and draws statistical power from both segment size, read-depth, and considers methylation call uncertainty. The count_dependency hypothesis setting will in a two-sample setting perform a Fisher exact test on binarized methylation call count, or with more samples, a χ^2^-test on the full contingency table. Most conservatively, the hypothesis bs_diff tests for a difference between mean read methylation rate between samples. Therefore, a methylation rate 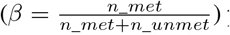 per read is computed on binarized methylation calls, and then a ranked test on methylation *β*-scores is performed. Thus, this test draws statistical power only from read-depth. Both the llr_diff and the bs_diff hypothesis perform a two-sided Mann-Whitney-U test in the two-sample case, and a two-sided Kruskal-Wallis test in the *n >* 2 sample case. Alternatively, if the parameter `–paired` is provided in a 2-sample setting, pycoMeth will instead compute *β*-scores for each genomic site observed in both samples and perform a two-sided Wilcoxon signed rank sum test on site-level *β*-scores.

As multiple testing correction on top of filtered results can introduce biases [37], pycoMeth Meth_Comp does not filter segments before p-value calculation safe for segments with insufficient data-points to compute the selected test. Multiple testing correction is thus performed across p-values from all segments with sufficient data, which leads to a large number of tests that need to be corrected for. Optionally, independent hypothesis weighting (IHW) [21] can be enabled to mitigate the problem of inflated p-values when testing a large number of segments for DMRs with p-value adjustment. When enabled, the scaled 1-centered standard deviation of methylation rates is used as a weight and multiplied with raw p-values. In any case, p-value adjustment is computed on raw or weighted p-values, and a large number of p-value adjustment methods are provided to users as implemented by the statsmodels python package [34].

### Methylome Segmentation Benchmark Setup

The pycometh methylome segmentation was compared to two existing tools designed for WGBS, methylKit [19] and MethCP [20]. MethylKit was used to perform a single-sample segmentation independent of differential methylation, wheres MethCP supports a 2-sample segmentation based on differential methylation. In order to evaluate the effect of segmentation granularity on DMR calling, we created a coarser segmentations with parameters `–window_size 600 –max_segments_per_window 16` (a maximum of 16 segments per 600 CpG calls) and a more fine-grained segmentation with parameters `–window_size 300 –max_segments_per_window 16` (a maximum of 16 segments per 300 CpG calls). No methylation rate prior was provided and haplotype information was provided as read-groups in the MetH5 format. For the segmentation using methylKit and MethCP, binarized metylation counts (LLR threshold 2.0) were created from the Nanopore methylation calls such as produced in a WGBS experiment. The methylKit segmentations were created based on total methylation rate of the compared samples/haplotypes. For MethCP methylation rates per sample/haplotype were computed. MethylKit segmentations were created using the function methSeg with parameters maxInt=100 and minSeg=10 as suggested in the methylKit documentation. MethCP was run with default parameters.

### Differential Methylation Testing setup

PycoMeth Meth_Comp was run in all of the three hypothesis options (bs_diff, llr_diff, count_dependency), with and without independent hypothesis weighing, and with p-value adjustment using the Benjamini-Hochberg method[38]. MethCP differential methylation testing was run with Fisher’s combined probability test. P-values reported by MethCP are already reported as adjusted by methylKit’s implementation of SLIM [39].

Intervals called by MethCP which were based on a single call were removed, since these obtained false significance from grouped Nanopolish calls being duplicated in the pseudo-bulk generation (**Supplementary Figure 2**)

## Acknowledgments

The present contribution is supported by the Helmholtz Association under the joint research school “HIDSS4Health - Helmholtz Information and Data Science School for Health”.

## Abbreviations

API: application programming interface
ASM: allele specific methylation
CGI: CpG-island
CLI: command line interface
CpG: 5’-cytosine-phosphate-guanine-3’
DMR: differentially methylated region
HDF5: Hierarchical Data Format (HDF) version 5
HMM: hidden Markov model
LLR: log-likelihood ratio
NCBI: National Center for Biotechnology Information
ONT: Oxford Nanopore Technologies

## Availability of data and materials

### Code availability

#### Software

- pycoMeth: https://github.com/snajder-r/pycometh
- meth5: https://github.com/snajder-r/MetH5Format

Benchmark scripts: https://github.com/snajder-r/benchmark_meth5

### Data availability

Sequencing raw data and variant calls were downloaded from the Genome in a Bottle (GIAB) consortium [22]. Download links are provided in **Supplementary Table 1**.

## Ethics approval and consent to participate

Not applicable

## Consent for publication

Not applicable

## Authors’ contributions

R.S. with guidance by M.B. and O.S. designed and developed the MetH5 format and Bayesian methylation segmentation method. A.L. and R.S. designed and implemented pycoMeth. R.S. prepared the figures and wrote the manuscript with input from M.B. and O.S.

## Additional Files

- **Supplementary Table 1**: `s1_giab_data.xlsx` -Benchmark data including download links

## Supplementary Materials

**Supplementary Figure 1:**
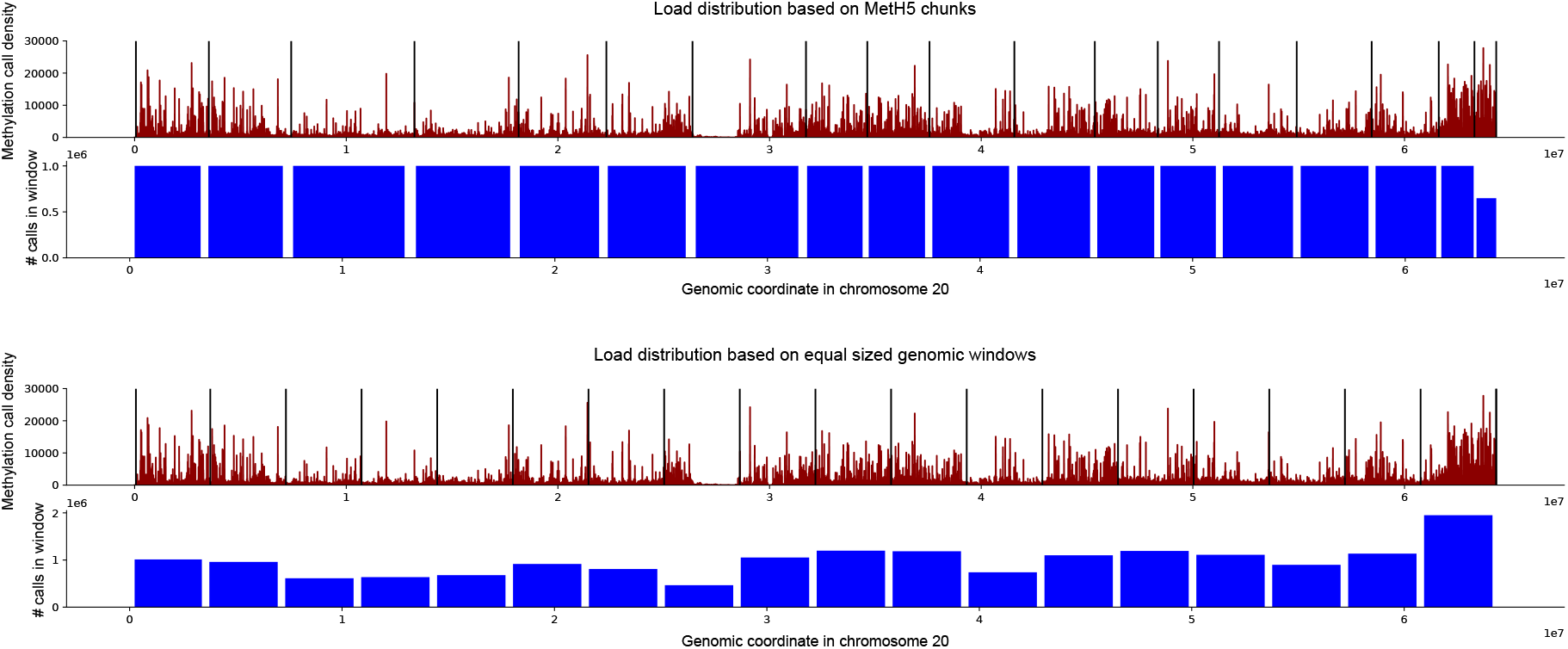
Parallel computing first requires splitting the input dataset for each parallel task. Good load distribution requires even distribution of input data. A naïve approach can be to distribute load based on evenly spaced genomic windows. However, CpG density and coverage are not even across an entire chromosome, leading to an uneven distribution of methylation calls per parallel task. MetH5 stores methylation calls in a chunked fashion. Definining parallel operations over MetH5 chunks results in an even load distribution across all parallel tasks. Methylation call density plotted here is computed as the total number (from all mapped reads) of methylation calls in a 1000bp window on HG002.

**Supplementary Figure 2:**
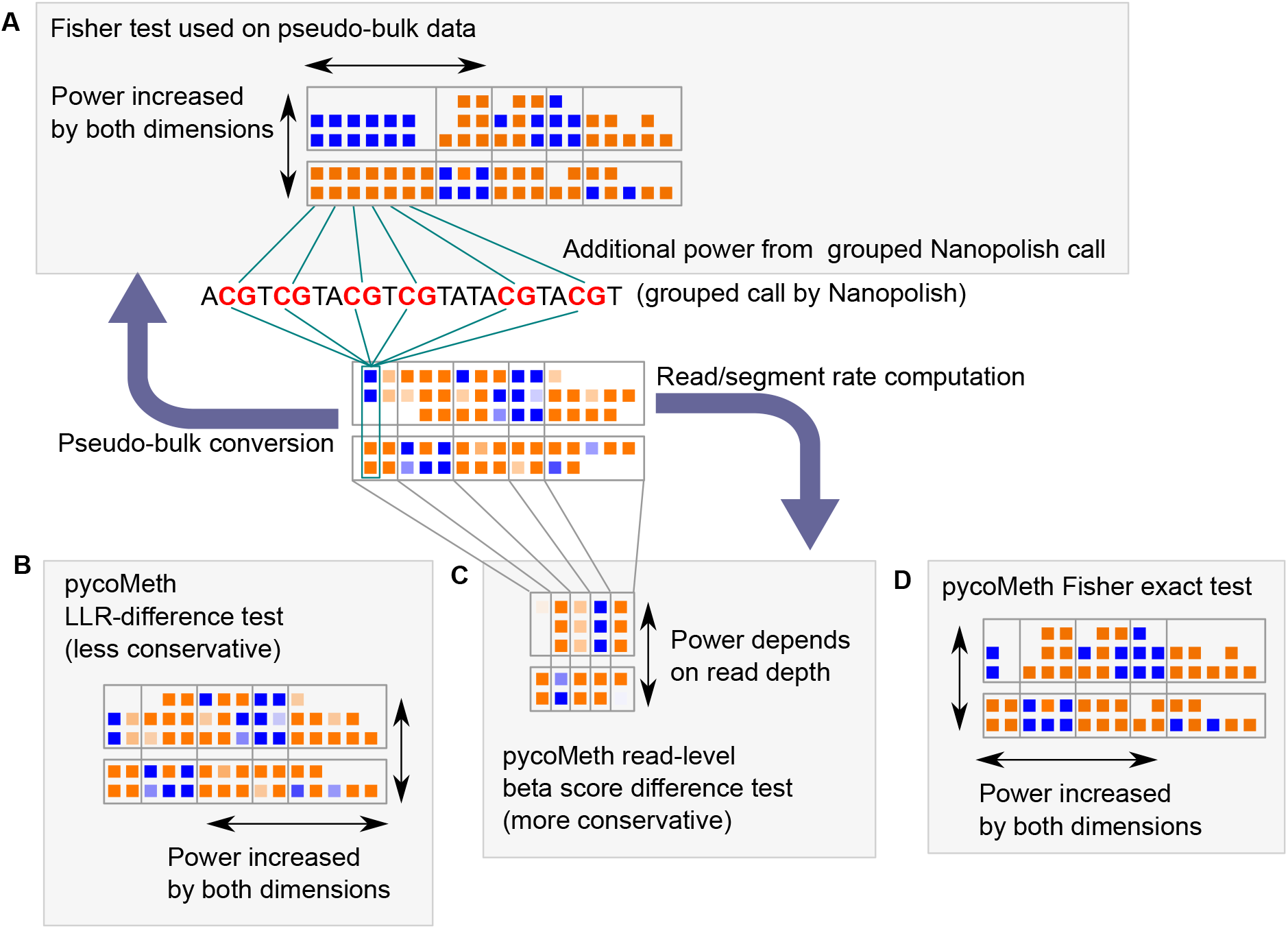
Illustration of where different differential methylation testing methods draw their power. **A**) In attempting to analyze Nanopolish methylation calls with tools developed for bulk bisulfite sequencing data, we create pseudo-bulk data. Tests generated for pseudo-bulk comparison (such as MethCP which we evaluated in this work) test based on CpG-level methylation rates and coverage across all reads and therefore draw power from the segment size and read depth. Furthermore, since Nanopolish generates grouped calls for nearby CpG-sites, some calls are therefore not independent and thus artificially generate more testing power. **B**) PycoMeth with the parameter “–hypothesis llr_diff” performs a less conservative test implemented in the pycoMeth package, where each individual methylation call is treated as independent and samples are compared based on their LLR distribution. Here discovery power is determined also by a combination of segment length and sequencing depth. **C**) PycoMeth with the parameter “–hypothesis bs_diff” instead computes a methylation rate per read per segment and draws power only from the independent information (sequencing depth). **D**) Fisher Exact test as implemented in pycoMeth for two-sample tests with the parameter “–hypothesis count_dependency”.

**Supplementary Figure 3:**
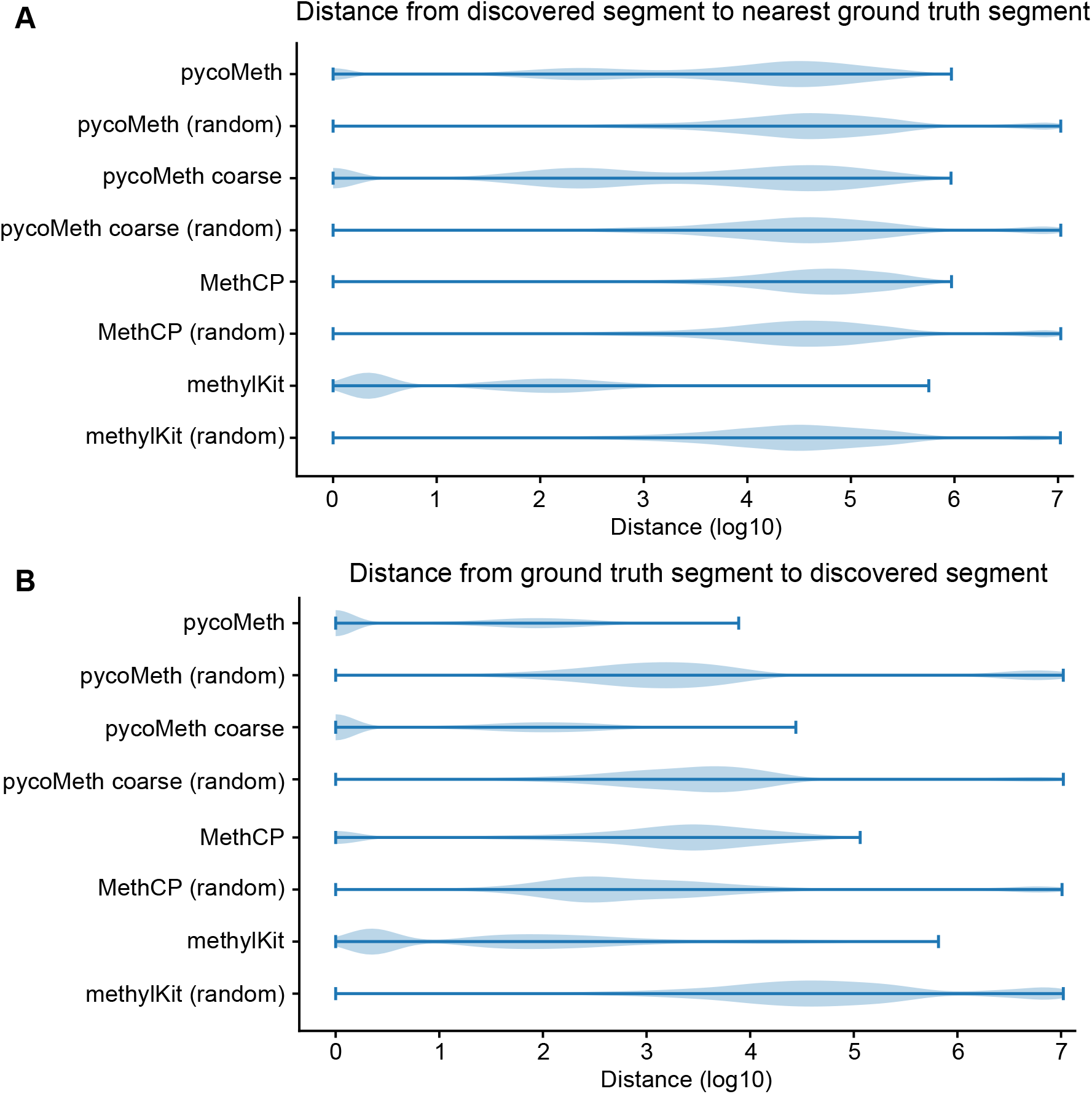
Permutation test on segmentations. In a permuted segmentation the original predicted segments retain their sizes but are shuffled in their order. This simulates a random segmentation with the same granularity. **A**) Distance from a discovered (predicted) changepoint to the nearest ground-truth changepoint. **B**) Distance from each ground-truth changepoint to the nearest predicted changepoint.

**Supplementary Figure 4:**
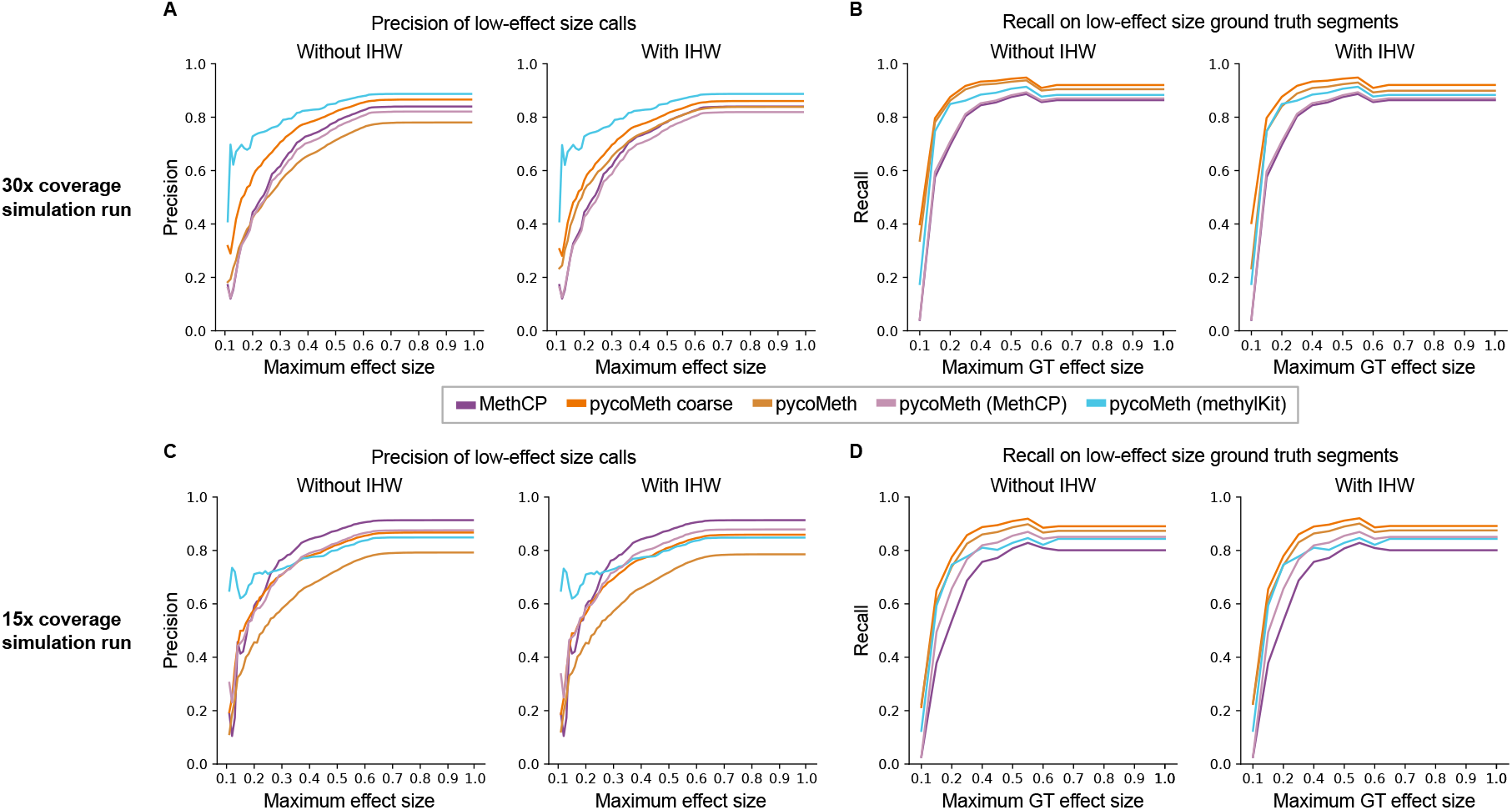
Investigating over-calling of low-effect size methylation calls. **A-B**) Precision and recall for effect-size capped DMR predictions with and without IHW on the high coverage simulation example. **C-D**) Matching analysis on the low coverage simulation example.

**Supplementary Figure 5:**
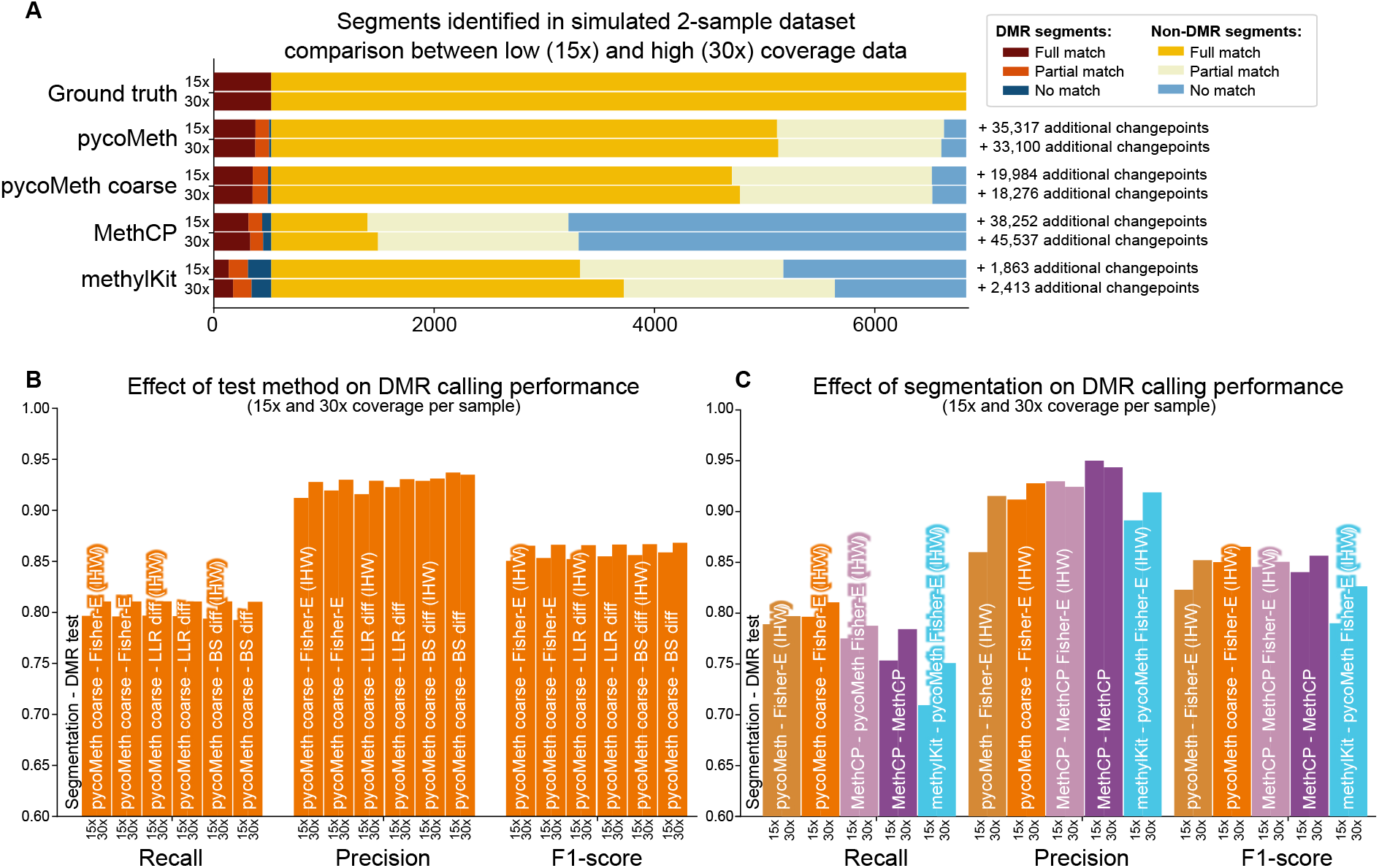
Benchmark on a simulated 2-sample dataset with lower coverage (15x). A) Compares the metrics analyzed in Figure 4A for 30x coverage simulated data with a lower coverage simulation of the same methylation profile at 15x. PycoMeth segmentations are largely unaffected, while MethCP and methylKit segmentations suffer from the drop in coverage. B) and C) show DMR calling recall, precision and F1-score for low coverage simulated data, analogous to Figure 4B-C.

**Supplementary Figure 6:**
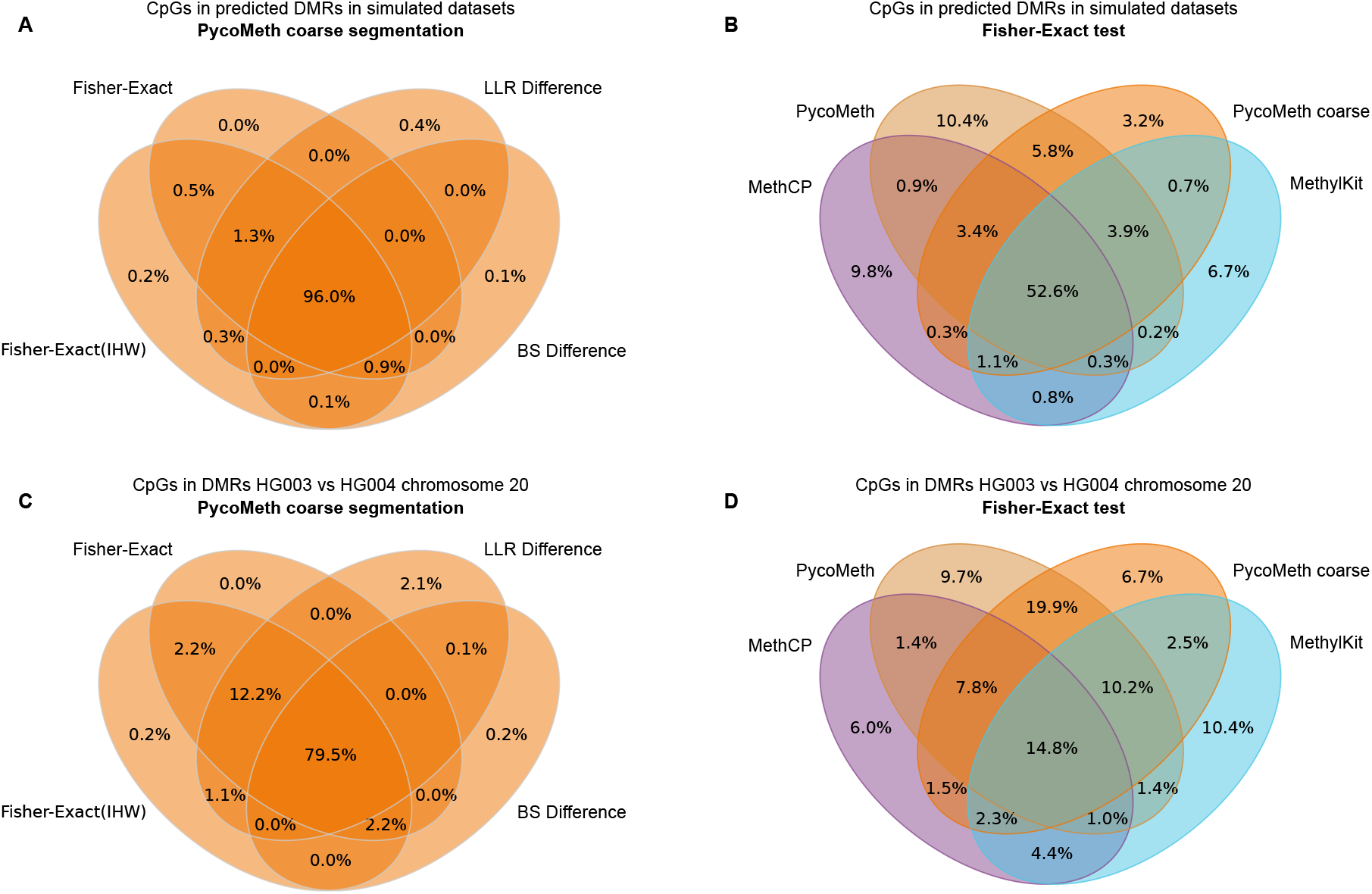
Agreement of different tests and segmentations measured as intersection of CpGs in DMRs called in the simulated data (**A-B**) and on chromosome 20 in the GIAB data parent comparison (**C-D**). Segmentation on the simulated data was overall more consistent, potentially due to the more homogeneous distribution of effect sizes.

**Supplementary Figure 7:**
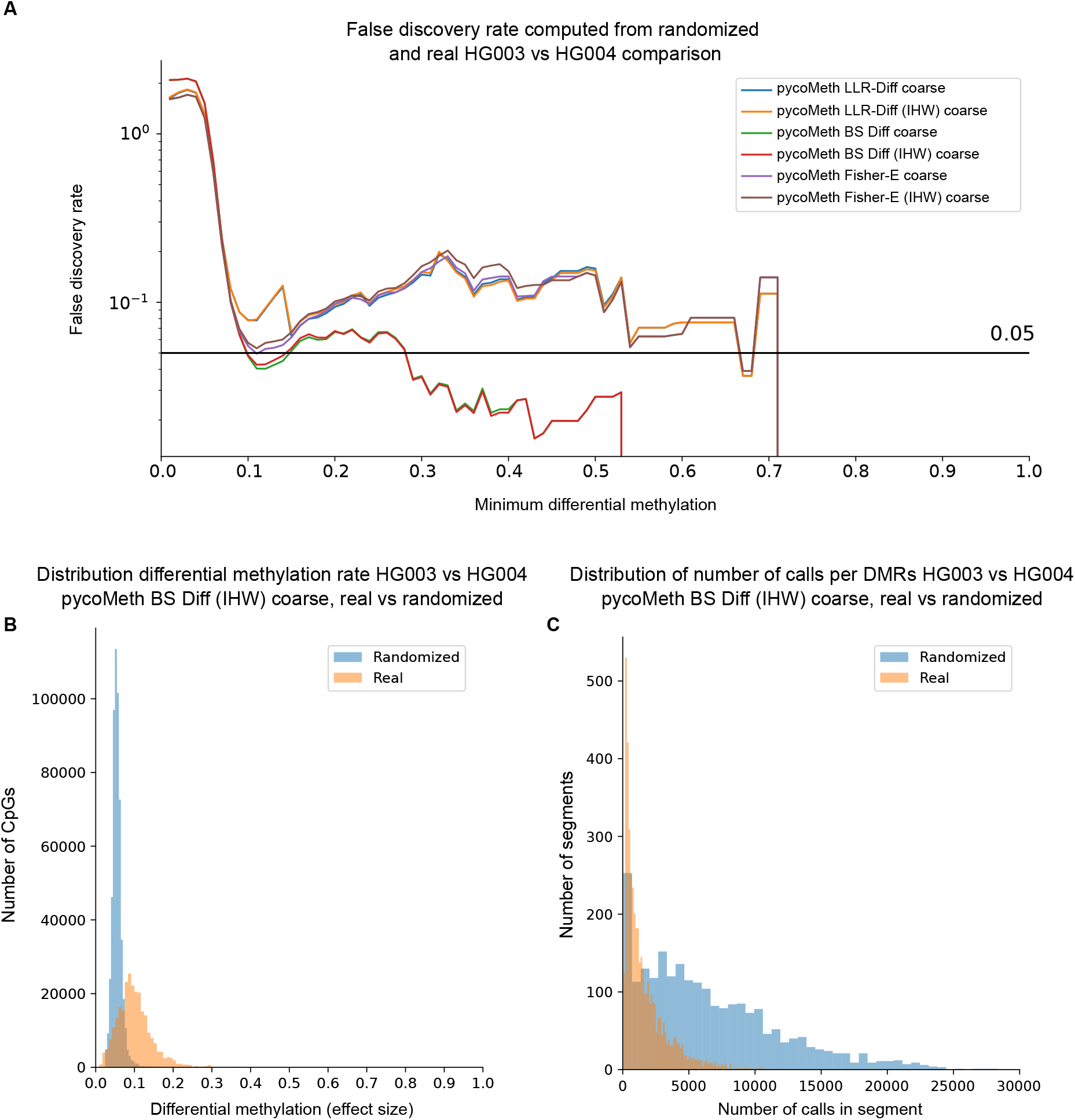
Randomization test results for comparison between HG003 and HG004. For the randomized dataset, for each sample and each chromosome LLRs have been shuffled to remove any read or site-dependent information. **A)** False discovery rate computed as the number of CpGs in DMRs in the randomized dataset divided by the same number for the real HG003 and HG004 comparison, plotted over the minimum segment differential methylation rate. We observe high FDR in segments with less than 0.1 differential methylation. The most conservative test implemented in pycoMeth (BS Diff) shows best FDR overall, and IHW appears to slightly reduce FDR in general. **B** Distribution of segment differential methylation (represented by CpGs in the segment) in called DMRs between the real and randomized dataset, from pycoMeth coarse with BS Diff test hypothesis and IHW. **C**) Distribution of calls per segment in called DMRs between the real and randomized dataset, from pycoMeth coarse with BS Diff test hypothesis and IHW.

## References

[1] Lisa D Moore, Thuc Le, and Guoping Fan. DNA methylation and its basic function. Neuropsychopharmacology, 38(1):23–38, January 2013.

[2] En Li and Yi Zhang. DNA methylation in mammals. Cold Spring Harb. Perspect. Biol., 6(5):a019133, May 2014.

[3] Suresh Kumar, Viswanathan Chinnusamy, and Trilochan Mohapatra. Epigenetics of modified DNA bases: 5-methylcytosine and beyond. Front. Genet., 9:640, December 2018.

[4] Sergey Kurdyukov and Martyn Bullock. DNA methylation analysis: Choosing the right method. Biology, 5(1), January 2016.

[5] Suhua Feng, Zhenhui Zhong, Ming Wang, and Steven E Jacobsen. Efficient and accurate determination of genome-wide DNA methylation patterns in arabidopsis thaliana with enzymatic methyl sequencing. Epigenetics Chromatin, 13(1):42, October 2020.

[6] Medhat Mahmoud, Nastassia Gobet, Diana Ivette Cruz-Dávalos, Ninon Mounier, Christophe Dessimoz, and Fritz J Sedlazeck. Structural variant calling: the long and the short of it. Genome Biol., 20(1):246, November 2019.

[7] Murray Patterson, Tobias Marschall, Nadia Pisanti, Leo van Iersel, Leen Stougie, Gunnar W Klau, and Alexander Schönhuth. WhatsHap: Weighted haplotype assembly for Future-Generation sequencing reads. J. Comput. Biol., 22(6):498–509, June 2015.

[8] Sergey Nurk, Sergey Koren, Arang Rhie, Mikko Rautiainen, Andrey V Bzikadze, Alla Mikheenko, Mitchell R Vollger, Nicolas Altemose, Lev Uralsky, Ariel Gershman, Sergey Aganezov, Savannah J Hoyt, Mark Diekhans, Glennis A Logsdon, Michael Alonge, Stylianos E Antonarakis, Matthew Borchers, Gerard G Bouffard, Shelise Y Brooks, Gina V Caldas, Haoyu Cheng, Chen-Shan Chin, William Chow, Leonardo G de Lima, Philip C Dishuck, Richard Durbin, Tatiana Dvorkina, Ian T Fiddes, Giulio Formenti, Robert S Fulton, Arkarachai Fungtammasan, Erik Garrison, Patrick G S Grady, Tina A Graves-Lindsay, Ira M Hall, Nancy F Hansen, Gabrielle A Hartley, Marina Haukness, Kerstin Howe, Michael W Hunkapiller, Chirag Jain, Miten Jain, Erich D Jarvis, Peter Kerpedjiev, Melanie Kirsche, Mikhail Kolmogorov, Jonas Korlach, Milinn Kremitzki, Heng Li, Valerie V Maduro, Tobias Marschall, Ann M McCartney, Jennifer McDaniel, Danny E Miller, James C Mullikin, Eugene W Myers, Nathan D Olson, Benedict Paten, Paul Peluso, Pavel A Pevzner, David Porubsky, Tamara Potapova, Evgeny I Rogaev, Jeffrey A Rosenfeld, Steven L Salzberg, Valerie A Schneider, Fritz J Sedlazeck, Kishwar Shafin, Colin J Shew, Alaina Shumate, Yumi Sims, Arian F A Smit, Daniela C Soto, Ivan Sović, Jessica M Storer, Aaron Streets, Beth A Sullivan, Françoise Thibaud-Nissen, James Torrance, Justin Wagner, Brian P Walenz, Aaron Wenger, Jonathan M D Wood, Chunlin Xiao, Stephanie M Yan, Alice C Young, Samantha Zarate, Urvashi Surti, Rajiv C McCoy, Megan Y Dennis, Ivan A Alexandrov, Jennifer L Gerton, Rachel J O’Neill, Winston Timp, Justin M Zook, Michael C Schatz, Evan E Eichler, Karen H Miga, and Adam M Phillippy. The complete sequence of a human genome. May 2021.

[9] Jared T Simpson, Rachael E Workman, P C Zuzarte, Matei David, L J Dursi, and Winston Timp. Detecting DNA cytosine methylation using nanopore sequencing. Nat. Methods, 14(4):407–410, April 2017.

[10] Shangqian Xie, Amy Wing-Sze Leung, Zhenxian Zheng, Dake Zhang, Chuanle Xiao, Ruibang Luo, Ming Luo, and Shoudong Zhang. Applications and potentials of nanopore sequencing in the (epi)genome and (epi)transcriptome era. Innovation (N Y), 2(4):100153, November 2021.

[11] Peng Ni, Neng Huang, Zhi Zhang, De-Peng Wang, Fan Liang, Yu Miao, Chuan-Le Xiao, Feng Luo, and Jianxin Wang. DeepSignal: detecting DNA methylation state from nanopore sequencing reads using deep-learning. Bioinformatics, April 2019.

[12] megalodon: Megalodon is a research command line tool to extract high accuracy modified base and sequence variant calls from raw nanopore reads by anchoring the information rich basecalling neural network output to a reference genome/transriptome,.

[13] Zaka Wing-Sze Yuen, Akanksha Srivastava, Runa Daniel, Dennis McNevin, Cameron Jack, and Eduardo Eyras. Systematic benchmarking of tools for CpG methylation detection from nanopore sequencing. Nat. Commun., 12 (1):3438, June 2021.

[14] Vladimir N Babenko, Irina V Chadaeva, and Yuriy L Orlov. Genomic landscape of CpG rich elements in human. BMC Evol. Biol., 17(Suppl 1):19, February 2017.

[15] Petr Danecek, James K Bonfield, Jennifer Liddle, John Marshall, Valeriu Ohan, Martin O Pollard, Andrew Whitwham, Thomas Keane, Shane A McCarthy, Robert M Davies, and Heng Li. Twelve years of SAMtools and BCFtools. Gigascience, 10(2), February 2021.

[16] Quincey Koziol and Dana Robinson. HDF5. [Computer Software] https://doi.org/10.11578/dc.20180330.1, March 2018.

[17] Helga Thorvaldsdóttir, James T Robinson, and Jill P Mesirov. Integrative genomics viewer (IGV): high-performance genomics data visualization and exploration. Brief. Bioinform., 14(2):178–192, March 2013.

[18] James T Robinson, Helga Thorvaldsdóttir, Wendy Winckler, Mitchell Guttman, Eric S Lander, Gad Getz, and Jill P Mesirov. Integrative genomics viewer. Nat. Biotechnol., 29(1):24–26, January 2011.

[19] Altuna Akalin, Matthias Kormaksson, Sheng Li, Francine E Garrett-Bakelman, Maria E Figueroa, Ari Melnick, and Christopher E Mason. methylkit: a comprehensive R package for the analysis of genome-wide DNA methylation profiles. Genome Biol., 13(10):R87, October 2012.

[20] Boying Gong and Elizabeth Purdom. MethCP: Differentially methylated region detection with change point models. J. Comput. Biol., 27(4):458–471, April 2020.

[21] Nikolaos Ignatiadis, Bernd Klaus, Judith B Zaugg, and Wolfgang Huber. Data-driven hypothesis weighting increases detection power in genome-scale multiple testing. Nat. Methods, 13(7):577–580, July 2016.

[22] Justin M Zook, David Catoe, Jennifer McDaniel, Lindsay Vang, Noah Spies, Arend Sidow, Ziming Weng, Yuling Liu, Christopher E Mason, Noah Alexander, Elizabeth Henaff, Alexa B R McIntyre, Dhruva Chandramohan, Feng Chen, Erich Jaeger, Ali Moshrefi, Khoa Pham, William Stedman, Tiffany Liang, Michael Saghbini, Zeljko Dzakula, Alex Hastie, Han Cao, Gintaras Deikus, Eric Schadt, Robert Sebra, Ali Bashir, Rebecca M Truty, Christopher C Chang, Natali Gulbahce, Keyan Zhao, Srinka Ghosh, Fiona Hyland, Yutao Fu, Mark Chaisson, Chunlin Xiao, Jonathan Trow, Stephen T Sherry, Alexander W Zaranek, Madeleine Ball, Jason Bobe, Preston Estep, George M Church, Patrick Marks, Sofia Kyriazopoulou-Panagiotopoulou, Grace X Y Zheng, Michael Schnall-Levin, Heather S Ordonez, Patrice A Mudivarti, Kristina Giorda, Ying Sheng, Karoline Bjarnesdatter Rypdal, and Marc Salit. Extensive sequencing of seven human genomes to characterize benchmark reference materials. Sci Data, 3:160025, June 2016.

[23] Nanopore community. https://nanoporetech.com/community,. Accessed: 2022-2-16.

[24] bonito: A PyTorch basecaller for oxford nanopore reads,.

[25] Justin M Zook, Jennifer McDaniel, Nathan D Olson, Justin Wagner, Hemang Parikh, Haynes Heaton, Sean A Irvine, Len Trigg, Rebecca Truty, Cory Y McLean, Francisco M De La Vega, Chunlin Xiao, Stephen Sherry, and Marc Salit. An open resource for accurately benchmarking small variant and reference calls. Nat. Biotechnol., 37 (5):561–566, May 2019.

[26] Heng Li. Minimap2: pairwise alignment for nucleotide sequences. Bioinformatics, 34(18):3094–3100, September 2018.

[27] modbampy. https://pypi.org/project/modbampy/,. Accessed: 2022-2-13.

[28] Ren-Hua Chung and Chen-Yu Kang. A multi-omics data simulator for complex disease studies and its application to evaluate multi-omics data analysis methods for disease classification. Gigascience, 8(5), May 2019.

[29] Andrew Collette. Python and HDF5. O’Reilly Media, Incorporated, 2013.

[30] Charles R Harris, K Jarrod Millman, Stéfan J van der Walt, Ralf Gommers, Pauli Virtanen, David Cournapeau, Eric Wieser, Julian Taylor, Sebastian Berg, Nathaniel J Smith, Robert Kern, Matti Picus, Stephan Hoyer, Marten H van Kerkwijk, Matthew Brett, Allan Haldane, Jaime Fernández Del Río, Mark Wiebe, Pearu Peterson, Pierre Gérard-Marchant, Kevin Sheppard, Tyler Reddy, Warren Weckesser, Hameer Abbasi, Christoph Gohlke, and Travis E Oliphant. Array programming with NumPy. Nature, 585(7825):357–362, September 2020.

[31] Pauli Virtanen, Ralf Gommers, Travis E Oliphant, Matt Haberland, Tyler Reddy, David Cournapeau, Evgeni Burovski, Pearu Peterson, Warren Weckesser, Jonathan Bright, Stéfan J van der Walt, Matthew Brett, Joshua Wilson, K Jarrod Millman, Nikolay Mayorov, Andrew R J Nelson, Eric Jones, Robert Kern, Eric Larson, C J Carey, ?Ilhan Polat, Yu Feng, Eric W Moore, Jake VanderPlas, Denis Laxalde, Josef Perktold, Robert Cimrman, Ian Henriksen, E A Quintero, Charles R Harris, Anne M Archibald, Antônio H Ribeiro, Fabian Pedregosa, Paul van Mulbregt, and SciPy 1.0 Contributors. SciPy 1.0: fundamental algorithms for scientific computing in python. Nat. Methods, 17(3):261–272, March 2020.

[32] McKinney. pandas: a foundational python library for data analysis and statistics. Python for high performance and scientific computing, 2011.

[33] Matthew D Shirley, Zhaorong Ma, Brent S Pedersen, and Sarah J Wheelan. Efficient “pythonic” access to FASTA files using pyfaidx. April 2015.

[34] Skipper Seabold and Josef Perktold. Statsmodels: Econometric and statistical modeling with python. In Proceedings of the 9th Python in Science Conference. SciPy, 2010.

[35] Plotly Technologies Inc. Collaborative data science. Montreal: Plotly Technologies Inc Montral, 2015.

[36] The Minh Luong, Vittorio Perduca, and Gregory Nuel. Hidden markov model applications in Change-Point analysis. December 2012.

[37] Maarten van Iterson, Judith M Boer, and Renée X Menezes. Filtering, FDR and power. BMC Bioinformatics, 11: 450, September 2010.

[38] Yoav Benjamini and Yosef Hochberg. Controlling the false discovery rate: A practical and powerful approach to multiple testing. J. R. Stat. Soc. Series B Stat. Methodol., 57(1):289–300, 1995.

[39] Hong-Qiang Wang, Lindsey K Tuominen, and Chung-Jui Tsai. SLIM: a sliding linear model for estimating the proportion of true null hypotheses in datasets with dependence structures. Bioinformatics, 27(2):225–231, January 2011.

